# Symbiosis reshapes the metabolism of sulfate-reducing bacteria in gutless marine worms

**DOI:** 10.1101/2024.12.09.627487

**Authors:** Grace D’Angelo, Manuel Kleiner, Anna Mankowski, Jerónimo Cifuentes-Anticevic, Marlene J. Violette, Valerie De Anda, Marc Mussmann, Eileen Kröber, Nicole Dubilier, Manuel Liebeke

## Abstract

Sulfate-reducing bacteria (SRB) are widespread in marine and terrestrial environments, where they often form syntrophic associations with bacteria, archaea, and eukaryotes. Among the most intimate of these are multipartite symbioses in gutless marine oligochaete worms, which host SRB and sulfur-oxidizing endosymbionts that engage in a syntrophic exchange of sulfur compounds. Despite decades of research on free-living SRB, the metabolic traits that enable SRB to persist in symbiosis, and how these differ across hosts and environments, remain poorly understood. Here, we show that a globally distributed clade of symbiotic SRB has a conserved core metabolism that diverges markedly from free-living relatives. Using comparative genomics and metaproteomics, we reveal that these endosymbionts retain key traits of SRB such as sulfate reduction, complete oxidation of acetate to CO_2_, amino acid degradation for nitrogen acquisition, and transport of essential nutrients. However, they exhibit a more oxygen-tolerant metabolism and lack typical nutrient-scavenging mechanisms of free-living SRB. One symbiont-specific trait, the glyoxylate bypass, was consistently expressed *in situ* and may serve both in reactive oxygen species defence and in biomass generation. The enrichment and expression of oxygen-tolerant pathways, coupled with the loss of nutrient-scavenging functions, indicate specialization to a host-associated, redox-fluctuating environment distinct from that of free-living SRB. Consistent with this shift, symbiont genomes are larger than those of free-living relatives, contrasting with genome reduction in many endosymbionts and reinforcing the importance of metabolic versatility. Our findings provide a framework for understanding how metabolic flexibility enables SRB to persist in long-term multipartite symbioses across diverse marine ecosystems.

## Introduction

Sulfate-reducing bacteria (SRB) are key drivers of carbon and sulfur cycling in anoxic environments, where they are often involved in syntrophic interactions with bacteria, archaea, and eukaryotes[1–10]. Some SRB occur transiently or facultatively on or within animal hosts; in gutless oligochaete worms, they are consistent and regular members of a multipartite symbiosis in a wide range of host species [11–17]. These small sediment-dwelling worms lack a mouth and gut, and live in an obligate nutritional symbiosis with up to 10 phylogenetically and metabolically diverse bacterial endosymbionts [12–15]. By migrating between oxidized and reduced sediment layers, the worms expose their symbionts to shifting redox conditions, allowing both their aerobic and anaerobic partners to remain metabolically active and coexist within the same host [12, 15].

Sulfate-reducing bacterial (SRB) symbionts occur in many gutless oligochaete species [17] alongside the primary endosymbiont of these animals: *Candidatus* Thiosymbion (hereafter *Thiosymbion*)[18]. *Thiosymbion* is a chemoautotroph that uses reduced sulfur compounds such as hydrogen sulfide as an electron donor to fix inorganic carbon into organic compounds [18, 19]. SRB symbionts and *Thiosymbion* exchange reduced and oxidized sulfur compounds in a syntrophic sulfur cycle thought to enhance protein yields and symbiotic productivity [12, 14, 15]. Despite a few detailed studies in mainly one host species [12, 15, 16, 20], the diversity and conservation of metabolic traits across hosts and their environments, as well as the extent to which symbiotic and free-living SRB differ, remain poorly understood.

To address these questions, we combined comparative genomics and metaproteomics to characterize a globally distributed clade of symbiotic SRB that occur in at least 14 gutless oligochaete species (17), previously referred to as “Delta 4” symbionts [16, 17, 21]. Our analyses revealed shared metabolic features and adaptations to a symbiotic lifestyle that distinguish these SRB from their free-living relatives. Given the monophyly of this globally distributed clade and its widespread, consistent association with oligochaete hosts and co-occurring symbionts, we propose the genus name “*Candidatus* Desulfoconcordia” [from de (L. “from”), sulfur (L. “sulfur”), and concordia (L. “union,” “harmony”)] for this clade.

## Results and Discussion

### *Desulfoconcordia* form a distinct symbiotic clade in the class Desulfobacteria

To investigate the evolutionary history of “*Ca.* Desulfoconcordia” (hereafter *Desulfoconcordia*), we compared a curated genomic dataset of cultivated and environmental SRB (**Table S1)** (21), with 44 metagenome assembled genomes (MAGs) of gutless oligochaete symbionts from 14 host species collected in coastal habitats around the world including Hawai’i, the eastern US, Australia, Italy, Belize, and the Bahamas [17] (**Table S2**). To cover a range of environments, hosts were collected from sediments around seagrass meadows and coral reefs with a few from sites with only sandy or fine-grained sediment (**Table S2**). Our phylogenomic analyses placed the *Desulfoconcordia* symbionts in a well-supported monophyletic clade within the class Desulfobacteria (**Fig. 1**, **Table S2**). Despite their broad geographic distribution, diverse habitats and varied host origins, all *Desulfoconcordia* symbionts clustered together, indicating a shared evolutionary history. Based on an average nucleotide identity ≥ 95% threshold for species delineation [23]), the *Desulfoconcordia* clade comprises at least 15 distinct species, with nine out of the 14 host species harboring a single, host-specific symbiont species (17, **Fig. S1**). The closest sequenced relatives of *Desulfoconcordia* symbionts included a MAG from a cellulose-fed sulfidic enrichment from the Black Sea (JAEINQ01) and several uncultivated SRB from hydrothermal vents, groundwater, and wastewater [22, 24, 25]. The nearest cultured representatives were members of the genera *Desulfonema*, *Desulfococcus*, and *Desulfosarcina* (**Fig. 1a**). The *Desulfoconcordia* symbiont genomes were on average 5.6 Mbp (**Fig. 1c**, **Table S2)**. In contrast, the closest free-living relatives of *Desulfoconcordia* had on average genomes of 4.1 Mbp (**Fig. 1c**, **Table S2**). Genome streamlining is typically observed in animal endosymbionts and arises as a consequence of an obligate host-associated lifestyle, resulting in progressive genome reduction [26]. In contrast, the expanded genomes of *Desulfoconcordia* compared to closely related free-living relatives may reflect selection for metabolic versatility required for life inside a motile host travelling through biogeochemically variable sediment layers.

**Figure 1:**
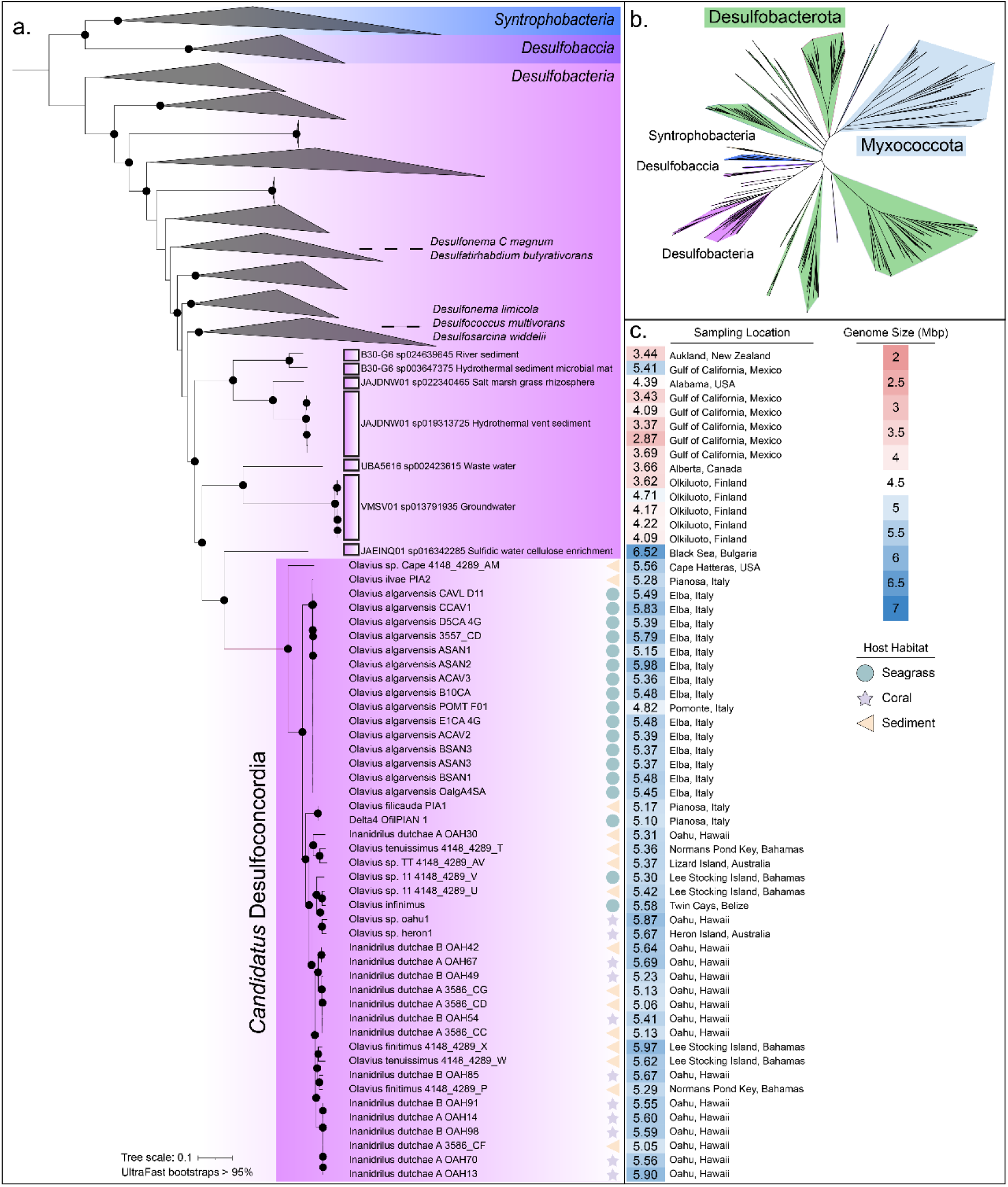
Phylogenomic placement of *Desulfoconcordia* symbionts. a) Phylogenomic tree based on 426 publicly available genomes and 44 *Desulfoconcordia* MAGs showing the placement of the symbionts within the class Desulfobacteria (purple/white box). Selected representative type strains are highlighted in their respective clades, and the closest relatives of the monophyletic *Desulfoconcordia* are labeled with their Genome Taxonomy Database (GTDB) genus-level identifiers (white box). Black dots show UltraFast bootstrap scores greater than 95%. b) Overview tree showing the placement of *Desulfoconcordia* within the phylum Desulfobacterota. c) Explanatory data showing genome size, sampling sites, and general habitat description. The trees were calculated using iqtree2 based on a) the GToTree workflow and a hidden Markov model (HMM) set of 74 bacterial single-copy genes and b) the marker gene sets assigned by checkM.

### A protein-based approach reveals a symbiont-specific protein signature

To identify metabolic features that distinguish *Desulfoconcordia* from free-living SRB, we applied the protein-based comparative approach previously used to analyse genomes from the phyla Desulfobacterota, SAR324, and Myxococcota [22, 27]. In this approach, genomes are hierarchically grouped based on the presence of absence of Pfam protein domains. This method defines groups of genomes with similar metabolic potential, here after metabolic guilds [28]. We compared the *Desulfoconcordia* genomes recovered in this study with those previously described in Langwig et al. using the same comparative framework [22](**Table S11**). In our extended analysis, the clusters (1-8) were largely consistent with those reported in the previous study (A-H) (**Fig. 2, Table S11**). Members of the class Desulfobacteria belonged to clusters F and D in the previous study and were distributed into clusters 6 and 7 here with a few in cluster 8. Unexpectedly, the *Desulfoconcordia* genomes formed a distinct metabolic guild (cluster 9) that was clearly separated from other SRB engaged in syntrophic or symbiotic relationships (in clusters 3, 6, and 7) (**Fig. 2**). Even though *Desulfoconcordia* are members of Desulfobacteria and share ecological similarities with other SRB symbionts, their genomes reveal a unique metabolic capability, placing them in a separate metabolic guild.

**Figure 2:**
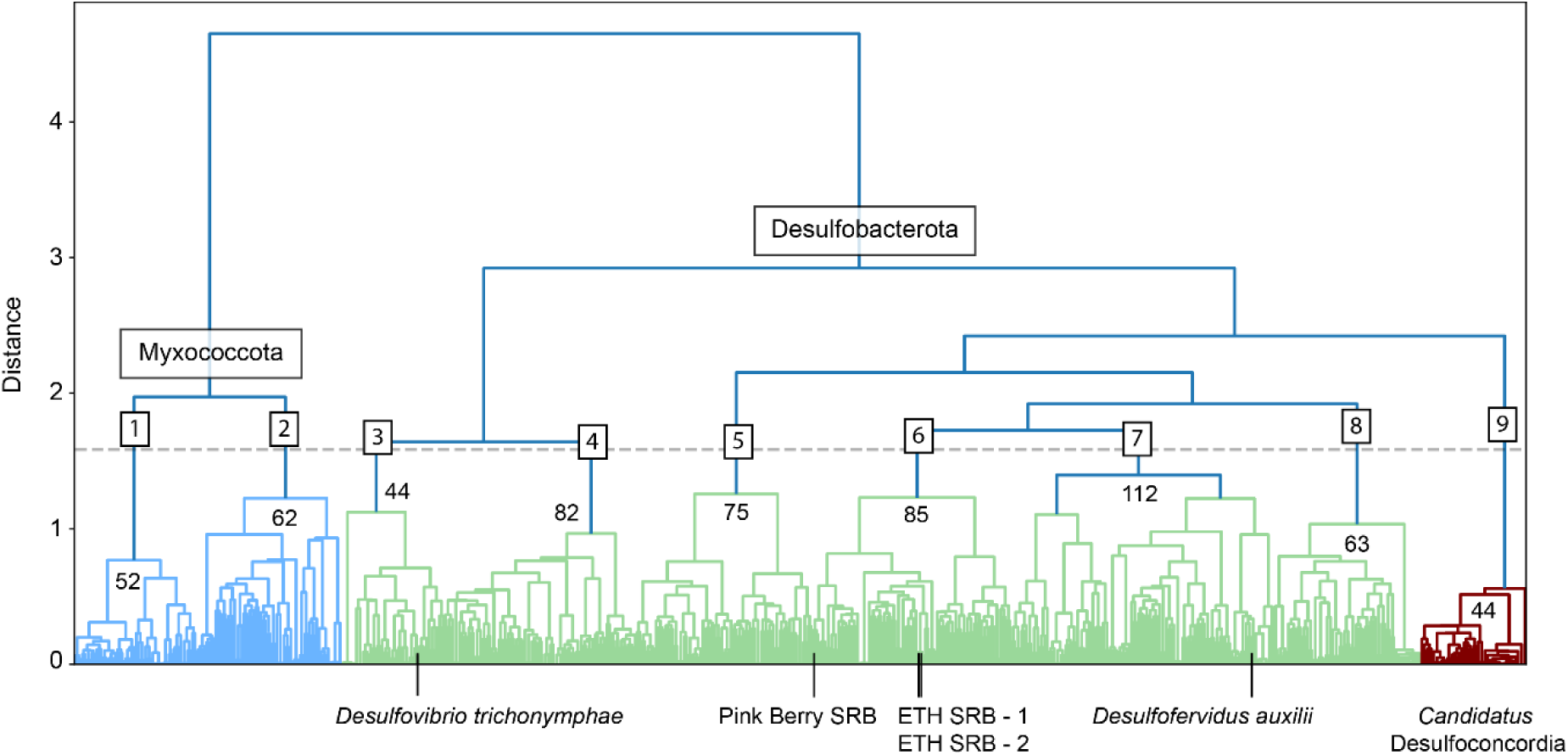
*Desulfoconcordia* have distinct protein domain profiles compared to other syntrophic Desulfobacteria. The dendrogram shows protein content similarity across 620 genomes, including 44 *Desulfoconcordia* MAGs, constructed using the Multigenomic Entropy Based Score (MEBS) tool (21, 26). Genomes were mapped against an existing Pfam-domain database and clustered based on their presence or absence using Jaccard distance and Ward variance minimization. The number of genomes in each cluster is indicated. Sulfate-reducing bacteria known to engage in syntrophic or symbiotic relationships are highlighted in their respective clusters.

To separate the metabolic features underlying this distinction, we analyzed the protein domain composition of the *Desulfoconcordia* metabolic guild. The *Desulfoconcordia* guild, when compared against the other clusters is characterized by *aceB* and *aceK* of the glyoxylate bypass, chorismate mutase, metallo-carboxypeptidases, amino acid oxidases, and domains of unknown function (**Table S8**). In contrast, domains present in other Desulfobacteria clusters (cluster 6, 7, and 8; **Fig. 2**) but absent from *Desulfoconcordia* genomes included the divalent ion tolerance protein CutA1 (cluster 6), Ppx/GppA phosphatase family (cluster 7), and PTS systems for sorbose/mannose uptake (cluster 8). Together, these differences define a symbiont-specific protein signature that distinguishes *Desulfoconcordia* from free-living SRB and points to adaptations related to their host-associated lifestyle.

### Comparative genomics identifies additional symbiotic traits of *Desulfoconcordia*

To further characterize the traits that distinguish *Desulfoconcordia* symbionts from their free-living relatives, we constructed a pangenome with symbiont genomes and their closest free-living relatives from GTDB. A total of 447 gene clusters comprised the core genome shared between the free-living and symbiotic bacteria, whereas 441 gene clusters were unique to the *Desulfoconcordia*, that is, shared by 100% of the symbiont genomes and absent from all free-living SRB **(Fig. 3a)**. These were defined as symbiont-only gene clusters (**Fig. 3a**)

**Figure 3.**
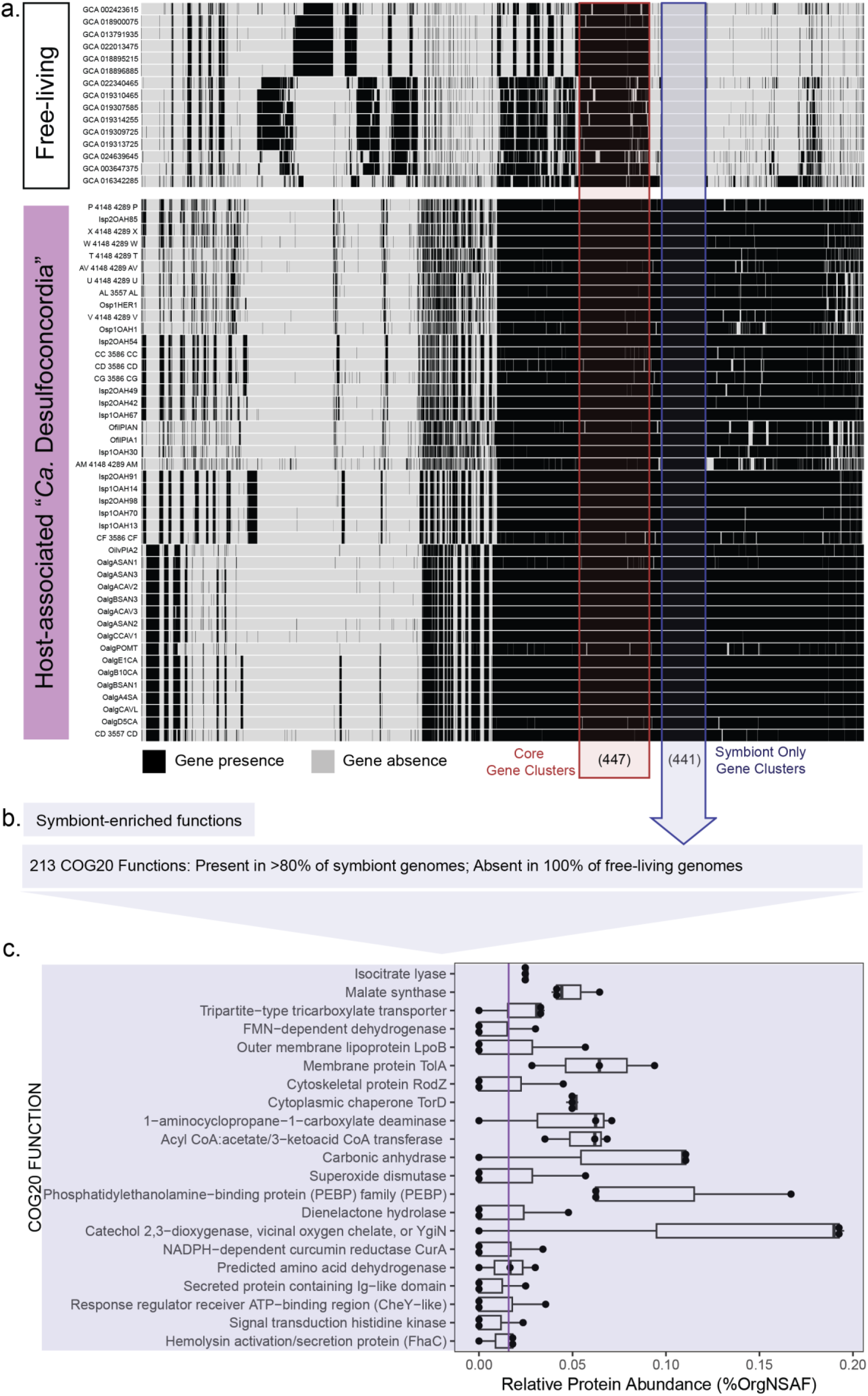
*Desulfoconcordia* are enriched in genes specific to their symbiotic lifestyle (a), and some of these were highly expressed in the metaproteomes of the gutless oligochaete *Olavius algarvensis* (c). a) A genomic comparison between *Desulfoconcordia* and related free-living SRB revealed 447 shared core gene clusters and 441 symbiont-specific gene clusters. X-axis labels are the genome accessions (**Table S2**). b) Functional enrichment analysis revealed 213 COG20 functional annotations present in at least 80% of symbiont genomes and absent from the free-living genomes. c) *Desulfoconcordia*-specific COG20 functions were expressed (19/213) in the host species *Olavius algarvensis* based on metaproteomic analyses. Metaproteomes were obtained from pooled samples (50 worms homogenized per sample, n=3). The median expression value is indicated with a vertical purple line. Relative protein abundance is presented as percent normalized spectral abundance factor (%NSAF) **(Tables S6 & S7)**.

To identify additional symbiont traits beyond the core genes, we searched for symbiont-enriched genes present in ≥ 80 % of symbiont genomes and absent from all free-living genomes. Using this threshold, a functional enrichment analysis revealed three KEGG modules and 213 COG20 functions that were enriched in *Desulfoconcordia* (**Fig. 3b, Tables S3 and S4**). The symbiont-enriched KEGG modules were the glyoxylate cycle, futalosine-dependent menaquinone biosynthesis, and pentose phosphate pathway, consistent with increased metabolic flexibility and redox balancing in a host-associated lifestyle. Among the 213 COG20 functions, 83 occurred in 100 % of the symbiont genomes, corresponding to highly conserved symbiotic traits. Examples include the two enzymes of the glyoxylate bypass, isocitrate lyase (AceA) and malate synthase (AceB), a Zn-dependent protease (YugP), benzoyl-CoA reductase, and transporters for amino acids such as ArgO and PutP (**Table S4**), suggesting an enhanced capacity for the use of diverse carbon and nitrogen sources.

Conversely, KEGG modules for fatty acid, lipopolysaccharide, and thiamine biosynthesis were present in 46%, 40%, and 33% of free-living genomes respectively, but absent from all symbiont genomes, consistent with reduced reliance on autonomous biosynthetic pathways in the host environment (**Table S3**). The TauABCD system, responsible for sulfonate transport, was also absent from all *Desulfoconcordia* symbionts, but present in 100% of the free-living SRB (**Table S4**). This system enables the use of aliphatic sulfonates such as taurine as sulfur sources when sulfate is scarce [29]. *Desulfoconcordia* symbionts have access to oxidized sulfur compounds from the surrounding sediment, as well as from the primary symbiont of gutless oligochaetes, *Thiosymbion*, which encodes genes for the oxidation of DMS to DMSO [30]. *Desulfoconcordia* can therefore rely on these abundant sulfur sources for both energy metabolism and the biosynthesis of sulfur-containing amino acids through cysteine synthase, rather than scavenging sulfur from taurine.

### *Desulfoconcordia* express symbiont-specific genes in the Mediterranean host *Olavius algarvensis*

Among gutless oligochaetes, *Olavius algarvensis* is the most comprehensively studied host species. Its symbiosis has been characterized in multiple genomic and ecological contexts, analyses that serve as the basis for linking *Desulfoconcordia* genomic traits to *in situ* protein expression [12–19, 21]. We therefore investigated the expressed metabolic pathways of the *Desulfoconcordia* symbiont in this host with metaproteomics. Host-derived proteins accounted for ∼75 % of total identifications and bacterial proteins for ∼25 % (**Table S5**). We identified over 800 *Desulfoconcordia* proteins, representing ∼3 % of the holobiont metaproteome (**Tables S5 and S6**). Using the symbiont-enriched gene sets defined in our comparative genomics analysis (**Fig. 3a and 3b**), we examined which of these traits were actively expressed *in situ*. We detected 19 of the 213 symbiont-enriched COG functions (**Table S7**) in the *Desulfoconcordia* proteomes (**Fig. 3c**).

The symbionts expressed the two enzymes of the glyoxylate bypass (isocitrate lyase (AceA) and malate synthase (AceB)), indicating that this function is not only enriched in the symbiont genomes but that this pathway operates as part of their regular metabolism in the natural environment. They also expressed genes involved in cellular integrity and stress protection, including cytoskeletal protein RodZ, superoxide dismutase, and carbonic anhydrase. By interconverting CO_2_ and bicarbonate, carbonic anhydrase can both stabilize intracellular pH under fluctuating redox conditions and maintain cellular CO_2_ concentrations for potential autotrophic fixation via the Wood-Ljungdahl pathway [31, 32]. Additionally, proteins associated with cell-cell interactions (e.g., secreted Ig-like domains and hemolysin-like secretion systems) suggest active communication between *Desulfoconcordia*, neighboring symbionts, and possibly the host epithelium. In other host-associated bacteria, hemolysins and related pore-forming proteins act as key mediators at the microbe-host interface, influencing adhesion, immune signaling, and phagolysosomal interactions [33]. These proteins in *Desulfoconcordia* may also contribute to adhesion and signaling at the host surface or participate in the controlled uptake and digestion of symbionts in phagolysosomes [20].

### Metaproteomics identifies the expressed nutrient and energy metabolism of *O. algarvensis Desulfoconcordia*

Beyond these symbiont-specific traits, the metaproteomic data enabled reconstruction of the core nutrient and energy metabolism of *Desulfoconcordia* in *O. algarvensis* (**Fig. 4, Fig. S2, Table S6**). *Desulfoconcordia* expressed transport systems for amino acids, peptides, and carbohydrates, together with core carbon oxidation pathways, indicating active exploitation of organic substrates available within the host environment. The diversity of ABC transporters for amino acids suggests reliance on uptake of host-available amino acids rather than *de novo* biosynthesis, potentially reducing energetic costs and reallocating resources toward growth and energy metabolism, as previously proposed [15]. Consistent with our comparative genomic analyses, pathways enriched in free-living SRB, including isoleucine, heptose, and thiamine biosynthesis and sulfonate scavenging via the TauABCD system, were not detected in either the *Desulfoconcordia* genomes or the metaproteomes of *O. algarvensis*. This supports the conclusion that these symbionts rely on host-associated nutrient sources rather than scavenging pathways typical of free-living relatives.

**Figure 4:**
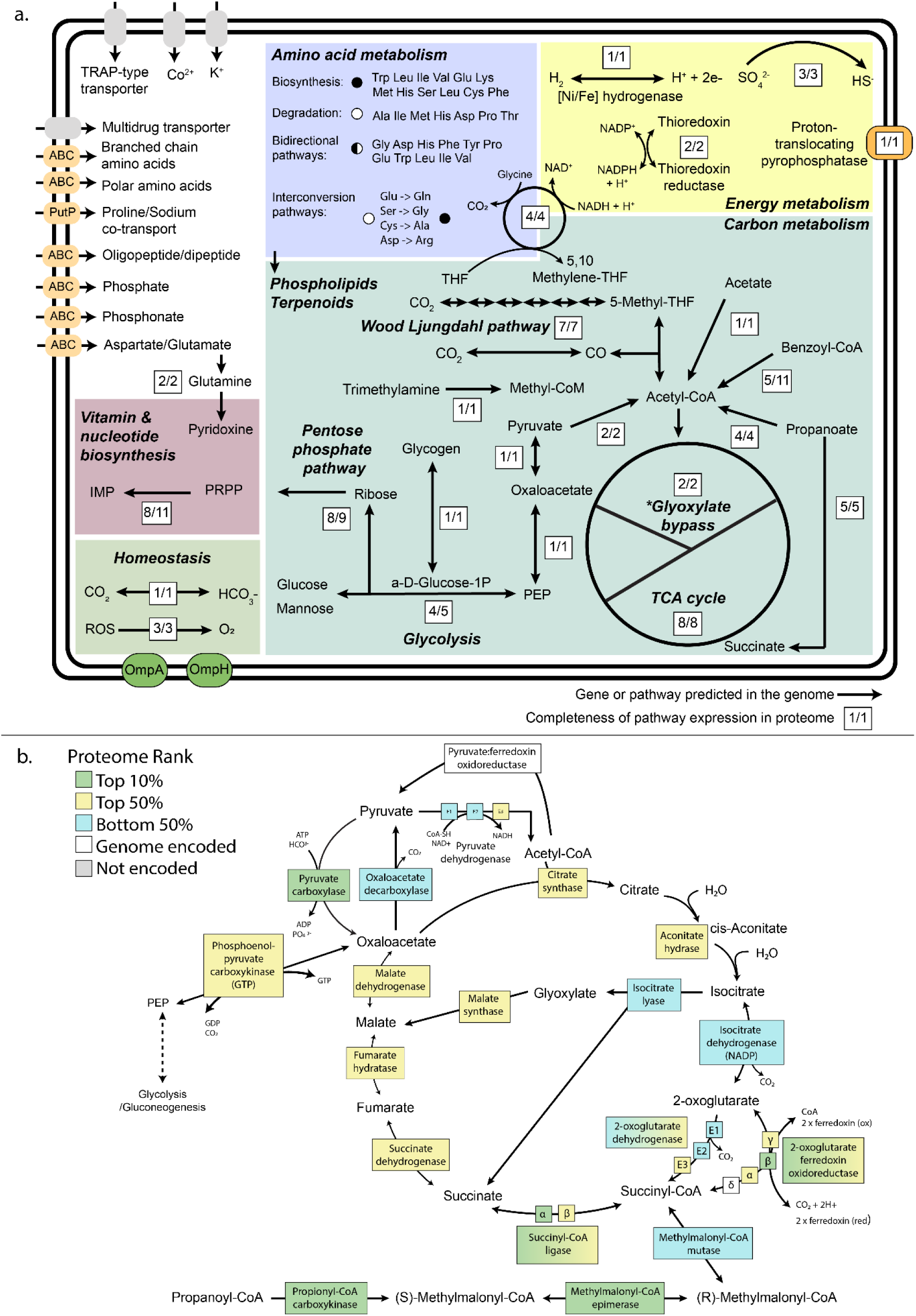
Expressed metabolic pathways in *Desulfoconcordia* from the host *Olavius algarvensis*. **a)** Arrows indicate the proteins and/or pathways detected in the metaproteome. The completeness of pathways is denoted by fractions. Amino acids (AA) are listed in one or more of the four AA metabolism groups if at least one marker gene for that AA was present, with black circles showing biosynthetic pathways, white circles degradation pathways and black/white circles bidirectional pathways. b) TCA cycle and glyoxylate bypass enzyme expression. Enzymes are colored according to rank in the metaproteome expression data (**Table S6**).

The symbionts also expressed the complete dissimilatory sulfate reduction pathway and the previously reported proton-translocating pyrophosphatase [16]. This enzyme hydrolyzes pyrophosphate (PPi), a by-product of sulfate reduction, and couples this reaction to proton translocation, thereby increasing ATP yield [16]. Recovery of energy from PPi likely enhances the bioenergetic efficiency of sulfate reduction, an advantage in the host-associated niche where energy availability may fluctuate with redox conditions during host migration.

Expression of both the methyl and carbonyl branches of the Wood–Ljungdahl pathway indicates metabolic flexibility that could operate in either direction-oxidatively for acetate catabolism or reductively for carbon fixation-depending on carbon and energy availability [15, 16, 34]. This bidirectionality would allow *Desulfoconcordia* to adjust carbon flux as environmental conditions shift between oxic and anoxic sediment layers.

The symbionts also expressed the glycine cleavage system, which catabolizes glycine, generates NADH for energy production, donates a methylene group to tetrahydrofolate, and releases ammonia as a nitrogen source [35]. This pathway may enable efficient recycling of nitrogen and carbon compounds, further contributing to metabolic efficiency.

Together, these expressed pathways reveal that *Desulfoconcordia* maintains a metabolically versatile physiology that enhances energetic efficiency, nutrient use, and redox resilience, supporting persistence within the chemically dynamic environment experienced during host migration.

### Clade-wide signatures of adaptation to fluctuating oxygen conditions

Although SRB are typically found in anaerobic environments, some are facultative anaerobes capable of tolerating or even exploiting transient oxygen exposure [8, 36, 37]. In free-living lifestyles and symbiotic associations, microbes often colonize microniches that match their redox capabilities [38–42]. In gutless oligochaetes, however, all symbiotic bacteria co-occur in an extracellular space just beneath the worm’s collagenous cuticle, which is permeable to gases and compounds of up to ∼70 kDa [12–14]. As a result, the symbiotic microbiome directly experiences the chemical environment of the surrounding sediment porewater, separated from it only by this thin permeable barrier. Because the host regularly migrates between oxic and anoxic sediment layers, the symbionts experience rapid and repeated changes in redox conditions (15). For the sulfate-reducing symbionts, these shifts require adaptations that either protect their oxygen-sensitive enzymes or provide alternative pathways for maintaining energy metabolism and redox balance.

To cope with shifting redox conditions, *Desulfoconcordia* encode and express the glyoxylate bypass, a modified branch of the tricarboxylic acid (TCA) cycle that supports both oxidative stress defense and carbon retention. The genes of the glyoxylate cycle (*aceA*, *aceB*, and the regulatory gene *aceK*) were present in 100% of *Desulfoconcordia* genomes and absent from their free-living representatives (**Fig. 3; Table S3**), and both AceA and AceB were expressed in situ (**Fig. 3c & 4b**). In addition, *Desulfoconcordia* expressed the 2-oxoglutarate:ferredoxin oxidoreductase, a reversible enzyme that is a typical marker of the reductive TCA (rTCA) cycle [43]. However, despite lacking Type II ATP citrate lyase and the complete canonical rTCA cycle [44], they can partially reverse the TCA cycle for anaplerotic carbon fixation and replenishment of biosynthetic intermediates via the glyoxylate bypass [43, 45].

Beyond this role in carbon metabolism, the glyoxylate bypass also contributes to oxidative stress tolerance: Glyoxylate can react directly with H_2_O_2_, providing a chemical sink for reactive oxygen species (ROS) generated during transient oxygen exposure [46]. The encoded and expressed isocitrate dehydrogenase is the NADPH-generating form, supplying reducing equivalents for ROS detoxification [47, 48]. Together, these features enable *Desulfoconcordia* to replenish TCA intermediates, support biosynthesis, and mitigate oxidative stress as the host migrates through oxygenated sediment layers.

### Conclusions

In this study, we identified and characterized the symbiotic clade *Desulfoconcordia* and demonstrated how their metabolism has been shaped by their host-associated niche. Using comparative genomics and metaproteomics, we identified symbiont-enriched traits, spanning central metabolism, stress tolerance, and host interaction, that distinguish them from their free-living relatives. The observed combination of redox-buffering capacities, central metabolic configurations, and symbiosis-associated functions illustrates that *Desulfoconcordia* can remain metabolically active and stably associated with their host despite its regular migration between oxic and anoxic sediment layers. Together, these results show how long-term host association in a dynamic redox environment can give rise to lineage-specific trait combinations. More broadly, this work advances our understanding of sulfur-based symbioses and highlights how metabolic and functional plasticity underpin stable microbial partnerships in marine ecosystems.

## Methods

### Sample collection

The metagenome-assembled genome bins of *Desulfoconcordia* from different host species from various field sites were generated in a previous study [17]. These genomes served as the basis for all metagenomic analyses in this study.

*Olavius algarvensis* worms for the density centrifugation metaproteomes were collected from the Sant’Andrea Bay on the island of Elba, Italy (7 m depth; 42°48’31.13”N, 10° 8’31.28”E) in September 2020. The animals were collected from the sediment by decantation with seawater and identified using both a dissection scope and fluorescence microscope. The density centrifugation protocol was carried out in the field and the samples were transported and stored at −80 °C prior to protein extraction and digestion.

### Metagenome collection

Raw metagenome sequences were published in previous studies: They are available in the European Nucleotide Archive (ENA) are under the Project IDs: PRJEB55913 and PRJEB94555 and those produced by DOE JGI are available under the GOLD Study ID Gs0095504. Metagenome assembled genomes (MAGs) were also published in parallel to PRJEB94555 in Zenodo under the DOI https://doi.org/10.5281/zenodo.17954011. MAGs were selected based on completeness and contamination values reported by CheckM[49]. All bins were over 80% complete and less than 10% contaminated. All 44 genomes used in this study had at least one rRNA operon and 18 unique tRNAs. *Desulfoconcordia* MAGs with genome sizes greater than 10% above the median genome size [5.6 Mbp] were manually refined in anvi’o[50, 51]. Read quality filtering, assembly, and binning were done as described in Mankowski *et al.,* https://github.com/amankowski/MG-processing_from-reads-to-bins/tree/main [17].

We imported the assemblies into anvi’o, and identified genes using Prodigal v2.6.3 in meta mode [52] We functionally annotated the genes against the COG24[53] and KOFam databases[54, 55]. KOfam annotation was improved following the adaptive adjustment of HMM thresholds implemented in *anvi-run-kegg-kofams* [56] Reads were recruited to the contigs using Bowtie2 v2.5.2[57] Binning results from the pipeline described in Makowski *et al.*[17], were imported to the anvi’o profile database. We identified the Desulfoconcordia bin by their GTDB annotation [58] obtained from running *anvi-run-scg-taxonomy*. We manually refined the *Desulfoconcordia* bins with *anvi-refine* based on their tetra-nucleotide frequency and coverage patterns [51].

### Phylogenomic analyses

We constructed two phylogenomic trees 1) to provide a broad overview of the Desulfobacterota phylum that was used in downstream protein functional domain analysis (**Fig. 1b**) and 2) a more specific look at the phylogenetic placement of the symbiont genomes in the Desulfobacteria class (**Fig. 1a, Table S1**). For the construction of the first tree, a set of previously curated genomes from NCBI [22] representing the Desulfobacterota and Myxococcota phyla were downloaded and dereplicated along with symbiont genomes using dRep (-pa 0.9 -sa 0.95 -comp 70 -con 50). The phylogenomic software iqtree2 was used to build a maximum likelihood tree (-m TEST -B 10000 --alrt 10000 -T AUTO) and the best fit model was determined to be LG+I+G4. To construct the second tree and determine the most well-supported outgroup based on all available sequences (rather than a pre-curated dataset), the GToTree workflow for building a phylogenomic tree with all available Genome Taxonomy Database (GTDB) reference sequences was used [59]. This second tree was constructed with the available GTDB sequences in the Desulfobacteria class with Syntrophobacteria and Desulfobaccia as outgroups. GToTree automatically identifies single-copy genes (SCGs) based on pre-curated hidden Markov model (HMM) gene sets, aligns with Muscle, trims with Trimal, builds a FastTree and can be piped directly into iqtree2 (https://github.com/AstrobioMike/GToTree/wiki/example-usage). The pre-curated Bacteria HMM set containing 76 gene targets was chosen to generate these trees. The FastTree step was followed by iqtree2 (-m TEST -B 10000 --alrt 10000 -T AUTO) with the best model fit determined to be Q.yeast+I+G4. The resulting trees were uploaded to iTol for visual annotation of taxonomy and for overlaying additional analyses [60].

### Pangenome Analysis

Anvi’o (v7.1) [50] was used to construct a pangenome that compares *Desulfoconcordia* symbionts to the other members of the well-supported (>95% bootstrap) GTDB free-living clade in which the symbiont genomes fall. The standard workflow for constructing a pangenome was followed (https://merenlab.org/2016/11/08/pangenomics-v2/). The contig databases were populated using all available hmm sources, KEGG module annotations, single-copy core gene taxonomy, tRNA scan, and the NCBI COG20 database (anvi-run-hmms, anvi-run-scg-taxonomy, anvi-scan-trnas, anvi-run-ncbi-cogs, and anvi-run-kegg-kofams). Functional enrichment analyses were performed between the “free-living” and “symbiotic” categories for both KEGG module completeness and COG20 functional annotations.

### Metabolic Annotation and Profiling

The MEBS software tool [61] was used to investigate and compare the metabolic capabilities of the assembled symbiont MAGs with respect to the reference genomes used in the first phylogenomic tree. The Prodigal gene predictions from the checkM output (generated by the dRep step described above) were used as the input for the MEBS scoring pipeline. Protein clustering was performed by first annotating the genomes against the most recently available Pfam database release (PfamV34) and then using the mebs_clust.py script to hierarchically cluster using Jaccard distance, Ward variance minimization, and a maximum distance threshold of 0.3. In accordance with the previous analysis, cluster distances were tested at 0.3, 0.35, 0.4, and 0.5 and 0.35 was chosen, as the clustering of the different classes in the Desulfobacterota and Myxococcota phyla closely resembled the result published in the initial study [22] **(Table S8, S9, S11)**.

### Metaproteomic analysis

Single *Olavius algarvensis* worms were pooled (n=50), homogenized and the symbionts were separated from each other and host debris by differential pelleting adapted from [62]. Homogenates were centrifuged for 2 minutes at 4 °C and 4000 x g (sample P1), the supernatant was transferred to a clean Eppendorf tube (sample S1), sterile filtered seawater (SFW) was added to sample P1; both P1 and S1 were centrifuged again with the same settings. The second supernatant from P1 was saved as S2 and again SFW was added to P1; S1 supernatant was transferred to a new tube (sample S3). P1 and S2 were centrifuged again for 2 minutes at 4 °C and 4000 x g. The resulting supernatant from P1 was discarded; the resulting supernatant from S2 was transferred to a new Eppendorf tube (sample S4). Samples S3 and S4 were centrifuged a final time for 7 minutes at 4 °C and 21000 x g and the resulting supernatant of S3 was saved in a new Eppendorf tube sample S5. Two of the resulting pellets (P1 most enriched in Cand. *Thiosymbion*, S3 most enriched in smaller symbionts (including Gamma3, Delta1, Delta4, Delta3, and the spirochete endosymbionts) and the final supernatant fraction (S5) were stored at −80 °C until further processing.

Tryptic peptides were prepared from the two pellet samples (P1, S3) and the supernatant (S5). The 1 ml of supernatant was lyophilized overnight prior to digestion. Protein was extracted and peptides generated following the filter-aided sample preparation (FASP) protocol adapted from Wiśniewski et al., [63]. 60 µl of SDT-lysis buffer (4% (wt/vol) SDS, 100 mM Tris-HCl (pH 7.6), 0.1 M dithiothreitol) was added to each sample and they were heated for 10 min at 95° C. The samples were centrifuged for 5 min at 21,000 x g and transferred to a 10-kDa molecular weight cutoff 500 µl centrifugal filter unit. Then, 400 µl of UA solution (8 M urea in 0.1 M Tris/HCl (pH 8.5)) was added to the filter units and the mixture was centrifuged for 40 min at 14,000 x g. Another 200 µl of UA was added and filter units were centrifuged again for 40 min at 14,000 x g. Samples were alkylated by adding 100 µl of IAA solution (0.05 M iodoacetamide in UA solution), mixing at 600 rpm for 1 min and incubating without mixing for 20 min at 22 °C in the dark. The filter units were centrifuged for 30 min at 14,000 x g and washed three times with 100 µl of UA by centrifugation for 20 min at 14,000 x g. The filters were then washed three times with 50 mM ammonium bicarbonate with the previous centrifuge settings. The filter units were transferred to fresh collection tubes and 0.8 ug of Pierce mass spectrometry MS-grade trypsin (Thermo Fisher Scientific) was added in 40 µl of ammonium bicarbonate buffer to each filter. The filters units were incubated overnight in a wet chamber at 37 °C. Peptides were eluted after the overnight incubation by centrifugation at 14,000 x g for 20 min. 50 µl of 0.5 M NaCl was added and mixed for 600 rpm for 1 min and the filter units were centrifuged again at 14,000 x g for 20 min. The eluted peptides were quantified using the Pierce micro bicinchoninic acid (microBCA) assay kit (Thermo Fisher Scientific), following the manufacturer instructions.

Peptide extracts were analyzed by on-line two-dimensional liquid chromatography coupled to mass spectrometry (2D-LC-MS/MS) using an Ultimate3000 RSLCnano system (Thermo Fisher Scientific) and Q-Exactive HF Orbitrap mass analyzer (Thermo Fisher Scientific) [64]. Samples were loaded onto a strong cation exchange (SCX) reverse-phase column (Thermo Fisher Scientific). During loading a C18 Acclaim PepMap 100 pre-column was inline downstream of the SCX column to capture peptides that did not bind to the SCX column. After loading, the downstream C18 Acclaim PepMap 100 pre-column was first placed in-line with the Easy-Spray PepMap C18 analytical column (75 µm by 75 cm; Thermo Fisher Scientific) and peptides were separated at 0.3 µL min^−1^ using a 120 min gradient from 95% buffer A (100% uHPLC water + 0.1% formic acid) to 31% buffer B (20/80 uHPLC water/100% acetonitrile + 0.1% formic acid) in 82 minutes, then to 50% B in 10 minutes, then to 99% B in 1 min, remaining at 99% B for 7 minutes, then dropping to 2% B in 1 minute and finishing at 98% buffer A / 2% buffer B. After this first elution, peptides were eluted from the SCX column to the C18 pre-column by injecting 20 µl of pH buffer plugs along a gradually increasing pH gradient (pH: 2.5, 3, 3.5, 4, 4.5, 5, 5.5, 6, 7, 8; CTIBiphase buffers, Column Technology, Inc.). After injection of each pH plug the C18 pre-column was again switched in-line with the C18 analytical column and the peptides were separated with the same elution gradient of A and B as stated above. This procedure resulted in 11 raw files per sample, each corresponding to the 100 min gradient described above. After each sample, the SCX column and C18 columns were each washed once (20 µl 4 M NaCl injected to SCX with a gradient of 100% eluent B at 0.3 µl min^−1^ and 20 µl injected to C18 with a gradient of 2% eluent B at 0.3 µl min^−1^) between samples. Eluting peptides were ionized via electrospray ionization (ESI) and analyzed in Q Exactive HF. Full scans were acquired in the Orbitrap at 60,000 resolution. The 15 most abundant precursor ions were selected in a data dependent manner, isolated with the quadrupole with a 1.2 m/z isolation window size, fragmented in the HCD cell with a NCE of 24, and measured in the Orbitrap at 15,000 resolution. Singly charged ions were excluded from MS/MS analysis. Dynamic exclusion was set to 25 s.

The mass spectrometry files were uploaded to Proteome Discoverer 2.5 and searched against a metaproteomic reference database (OlaviusV13.fasta). This custom protein sequence database was generated using a previously published *O. algarvensis* symbiosis database (PRIDE accession number PXD007510). The database was refined using CD-HIT to cluster highly similar sequences and by adding symbiont and host sequences with improved annotations [65]. The final database comprised 1,439,552 protein sequences including symbiont and host proteins and the common Repository of Adventitious Proteins (cRAP http://www.thegpm.org/crap) to represent common laboratory contaminants. MS/MS spectra were searched against the database using Proteome Discoverer vs. 2.5 (Thermo Fisher Scientific). The Precursor Detector node was set with an S/N threshold of 1.5 and the Sequest HT node search parameters included trypsin (full), a maximum of two missed cleavages, and a fragment mass tolerance of 0.1 Da. Dynamic modifications included acetylation of protein N termini (+42.011 Da), oxidation of methionine (+ 15.995 Da), and deamidation (+0.984 Da) of asparagine, glutamine, and arginine, with a maximum of 3 equal dynamic modifications per peptide. The static modification was carbamidomethylation of cysteine (+ 57.021 Da). False discovery rates (FDRs) for peptide spectral matches (PSMs) were calculated and filtered using the Percolator Node in Proteome Discoverer. Percolator was run with a maximum delta Cn 0.05, a strict target FDR of 0.01, and a relaxed target FDR of 0.05 and validation based on q-value. The Protein FDR Validator Node in Proteome Discoverer was used to calculate q-values for inferred proteins based on the results from a search against a target-decoy database. Proteins with a q-value of <0.01 were categorized as high-confidence identifications and proteins with a q-value of <0.05 were categorized as medium-confidence identifications. We combined all search results into a multiconsensus report and proteins were filtered to retain only those with an overall FDR of 0.05 or less. Normalized spectral abundance factors (NSAFs) were calculated for each bacterial and animal species (OrgNSAF) and multiplied by 100 to represent relative protein abundance as a percentage.

## Author contributions

GD, ML, and ND conceived the study. GD collected metagenomic data and performed phylogenomic, metabolic annotations, and pangenome analyses. AM generated the “*Ca*. Desulfoconcordia” metagenome-assembled genomes (MAGs). JCA performed MAG refinements. JCA and VDA provided input on the phylogenomics and MEBS analyses respectively. GD, MV, and MK generated mass spectrometry data and performed metaproteome analyses. MK provided conceptual input to the metaproteomic experimental design and data analysis. MM provided input on the metabolic reconstructions. GD, EK, ML, and ND wrote the manuscript with contributions from all authors.

## Acknowledgements

We thank Silke Wetzel (Max Planck Institute for Marine Microbiology) for help with laboratory work for this project. We thank Miriam Weber and Christian Lott (HYDRA Marine Sciences GmbH) for their support collecting samples in the field and Olivier Lemaire (Max Planck Institute for Marine Microbiology) for helpful discussions. All LC-MS/MS measurements were made in the Molecular Education, Technology, and Research Innovation Center (METRIC) at North Carolina State University. This study was funded by the Max Planck Society, the Gottfried Wilhelm Leibniz-Prize of the German Research Foundation (DFG) to Nicole Dubilier, and the US National Science Foundation (grant IOS #2426305 to Manuel Kleiner). The LLM ChatGPT was used to correct written text as a spelling and grammar check.

## Data availability

Previously published MAGs are available under https://doi.org/10.5281/zenodo.17954011. Additional MAGs refined and used in this study are available under https://doi.org/10.5281/zenodo.18889475 Proteomics data that were reanalyzed are available in the PRIDE repository under the Project ID: PXD014591 and proteomics data generated for this study will be available under Project ID: PXD075566 upon acceptance to a journal.

## Notes

### Competing Interest Statement

The authors have declared no competing interest.

### Summary of Updates

Author added and author affiliations updated. Figures 1 - 4 revised. Results text associated with figures updated to reflect the figure changes. Supplement files updated.

