## supplement_text for "Symbiosis reshapes the metabolism of sulfate-reducing bacteria in gutless marine worms"

### Supplementary text

**The core metabolism of the *Desulfoconcordia* symbionts is stable across host individuals**

To complement the bulk metaproteomics approach, we reanalyzed published protein expression data from individual worms. While the metaproteomes generated from 50 worms allowed us to target low-abundance symbiont proteins, we also investigated potential variations in gene expression at the individual host level that might be masked by a bulk approach (**Fig. S3**). Overall, the top abundant proteins in *Desulfoconcordia* were similar across individual worms (**Table S10**, **Fig. S3**) and consistent with those in the bulk analysis (**Table S6**). In all *Desulfoconcordia* proteomes from single-host metaproteomes, the most highly expressed gene was a hypothetical protein, sharing approximately 50% sequence identity (e-value of 10^-109^) with porin family proteins from members of the Desulfobacterota phylum in the NCBI BLAST database. Among the top 10 most abundant proteins were ABC transporters for amino acids and GroEL, a chaperone protein. Additionally, the anaerobic bidirectional form of carbon monoxide dehydrogenase of the Wood-Ljungdahl pathway was consistently within the top 100 proteins of single-host individuals. These complementary analyses provide a comprehensive understanding of *Desulfoconcordia* in *Olavius algarvensis* and a more detailed view of the variations in gene expression at the individual community level (**Fig. S3**).

**Reanalysis of published data**

Publicly available metaproteomic data were downloaded from the PRIDE (https://www.ebi.ac.uk/pride/) repository for reanalysis (Project ID: PXD014591)[1]. These mass spectrometry files were searched in Proteome Discoverer 2.5 against an updated metaproteomic reference database (OlaviusV13.fasta). The MS/MS spectra were searched against this reference database using the Sequest HT algorithm in Proteome Discoverer 2.5 (Thermo Fisher Scientific) with the following parameters: trypsin (full), maximum of two missed cleavages, 10 pp precursor mass tolerance, 0.1-Da fragment mass tolerance, and a maximum of three dynamic modifications per peptide. False discovery rates (FDRs) for peptide spectral matches (PSMs) were calculated and filtered using the Percolator node in the Proteome Discoverer. The Protein FDR Validator node was used to calculate q values for proteins based on the false discovery rates calculated by searching against the target-decoy database. Proteins with q values of < 0.01 were categorized as high-confidence and those with q-values between 0.01 – 0.05 were categorized as medium-confidence. We combined all search results into a multiconsensus report and proteins were filtered to retain only those with an overall FDR of 5% or less. Normalized spectral abundance factors (NSAFs) were calculated for each bacterial and animal species (OrgNSAF) and multiplied by 100 to represent relative protein abundance as a percentage.

1.	Jensen M, Wippler J, Kleiner M. Evaluation of RNAlater as a field-compatible preservation method for metaproteomic analyses of bacterium-animal symbioses. *Microbiol Spectr* 2021;**9**:e0142921. https://doi.org/10.1128/Spectrum.01429-21

### Supplement Figure Legends

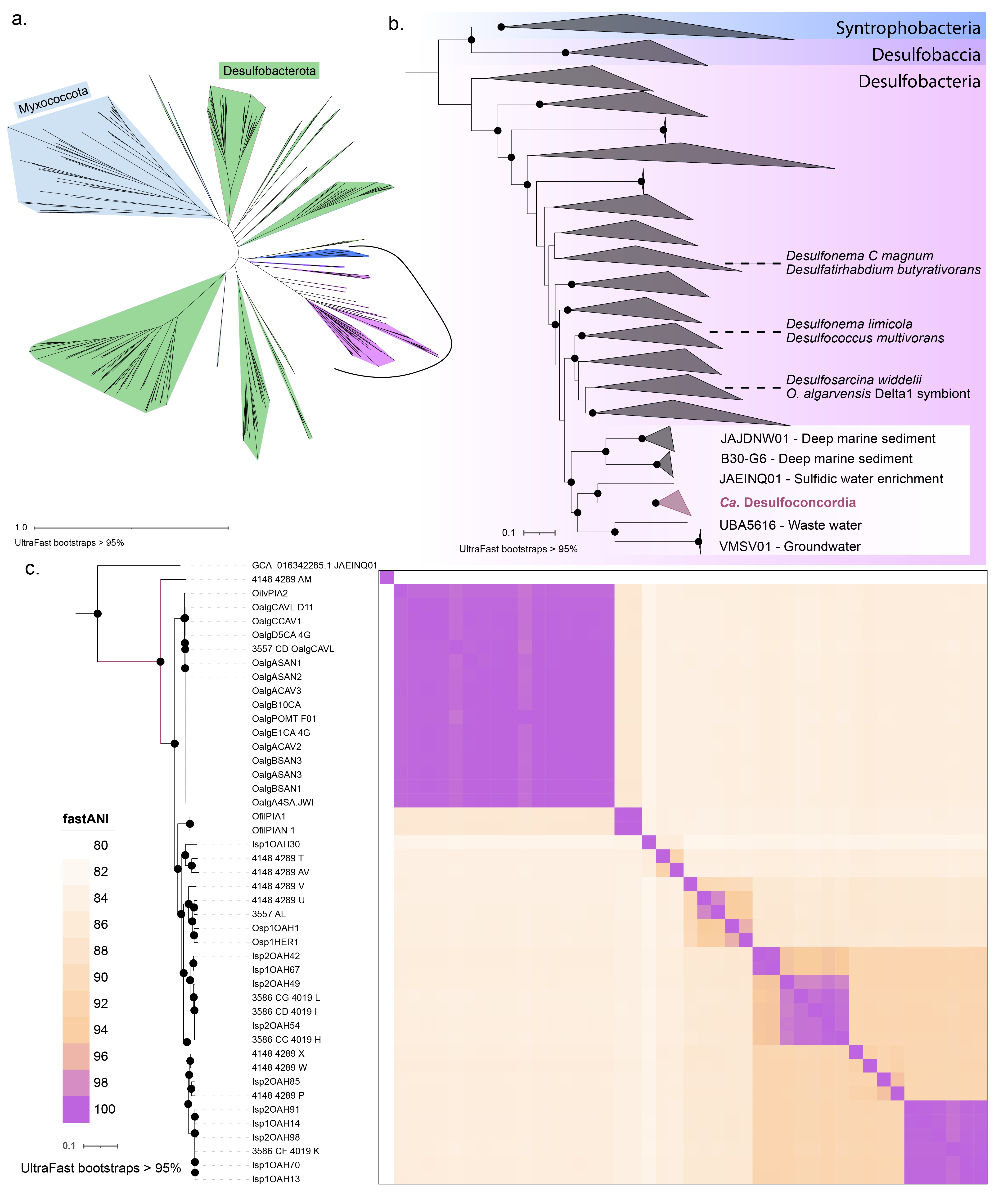

**Figure S1: Phylogenomic placement of *Desulfoconcordia* symbionts.** The *Desulfoconcordia* symbiont MAGs are displayed in their phylogenomic context according to the GToTree workflow for GTDB reference genomes. a) Phylogenomic tree generated with 426 publicly available genomes and 14 *Desulfoconcordia* MAGs places these symbionts in the Desulfobacteria class. b) For a more detailed view, the classes Syntrophobacteria and Desulfobaccia were used as outgroups for the class Desulfobacteria. Selected representative type strain members of Desulfobacteria are highlighted in their respective clades. Close representatives to the monophyletic *Desulfoconcordia* symbiont clade are labeled with their GTDB taxonomic genus-level identifiers. Black dots represent UltraFast bootstrap scores greater than 95%. c) Average nucleotide identity scores calculated using fastANI with a color scale delineation representing the 95% cutoff for species distinction.

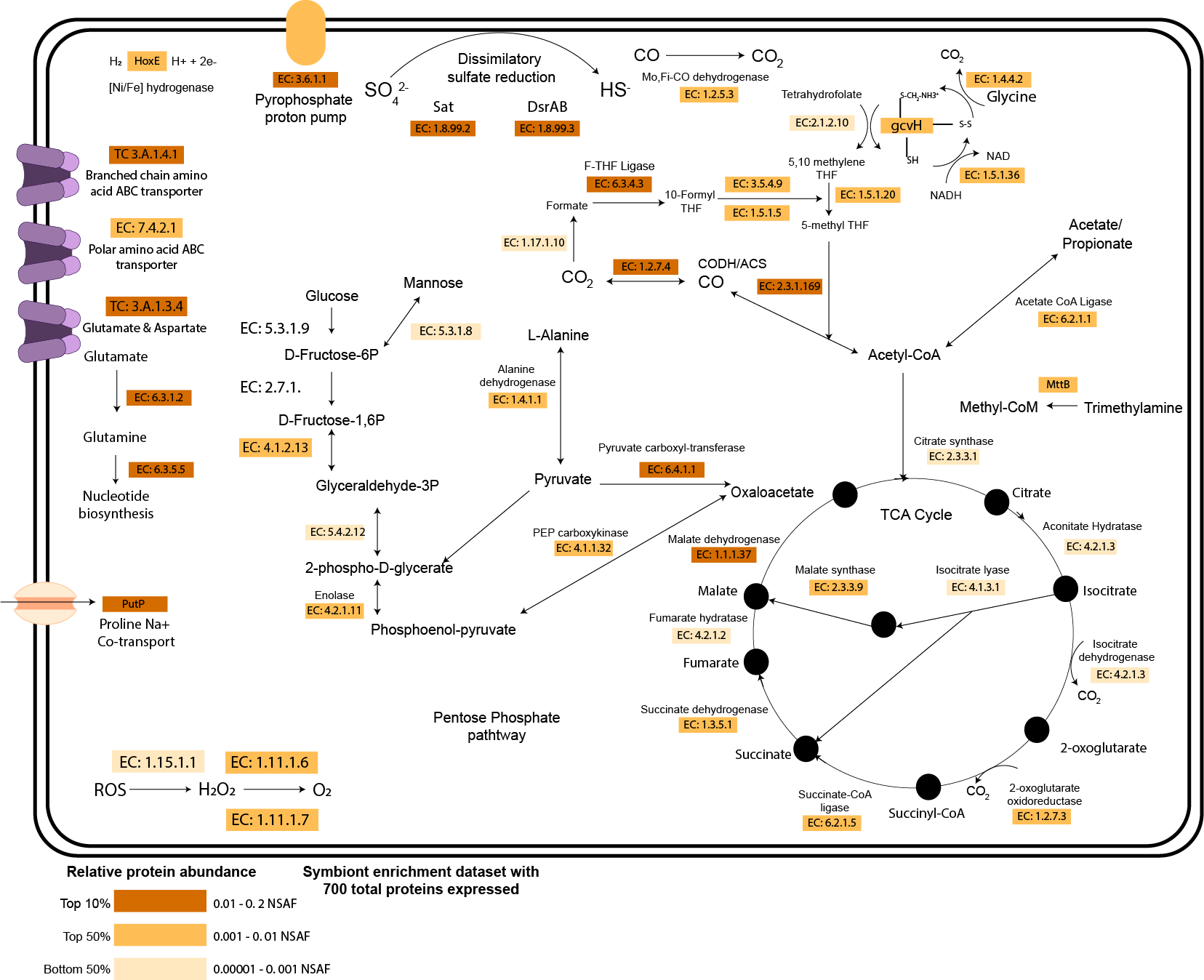

**Figure S2: Detailed metabolic overview of *Desulfoconcordia*.** A metabolic reconstruction of the pathways from the simplified metabolic overview based on enrichment samples (n=50 worms homogenized). EC numbers or gene names of key enzymes in each pathway are highlighted in different shades of orange to represent their respective relative abundance within the metaproteome.

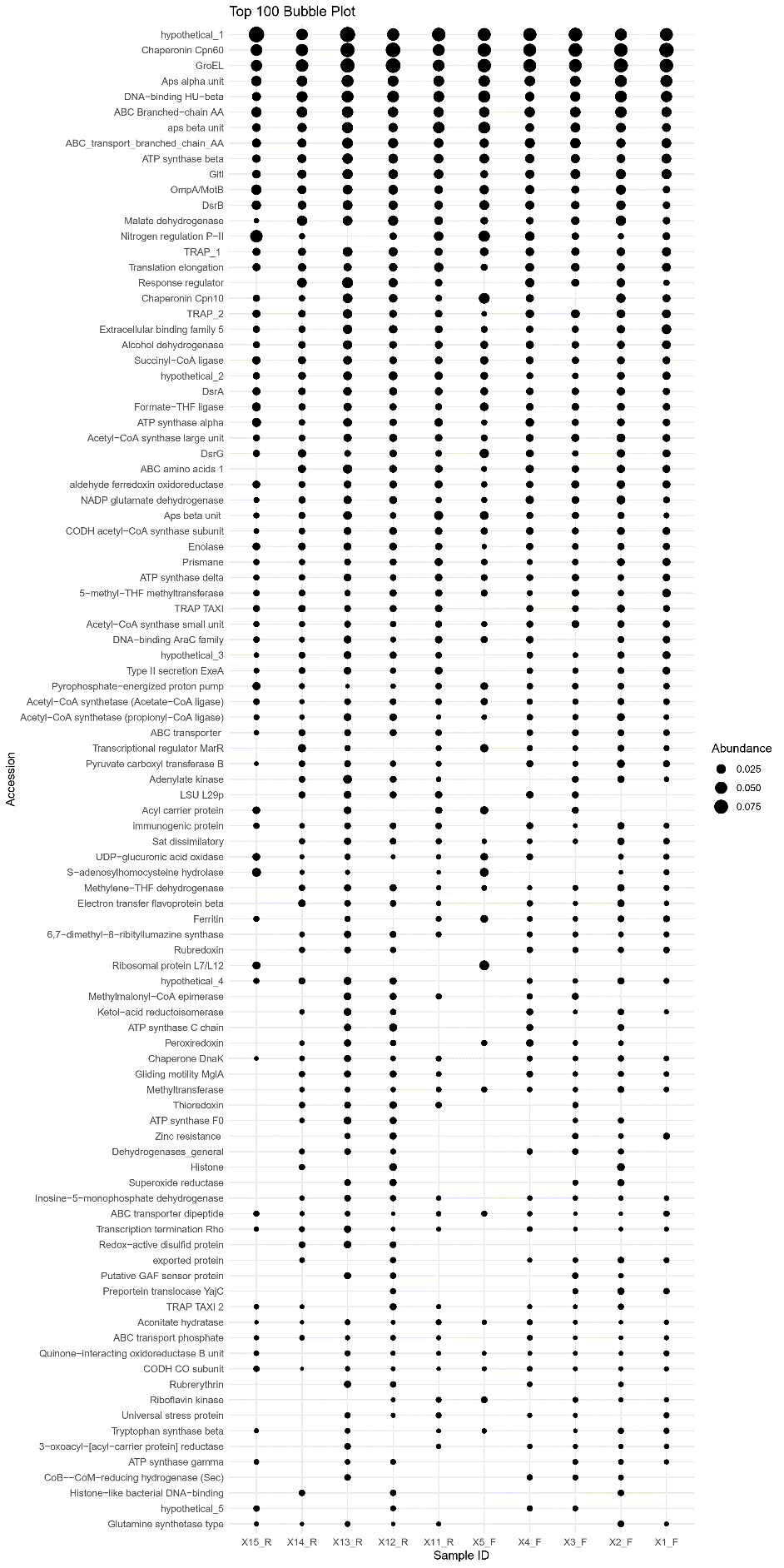
**Figure S3. Top 100 expressed *Desulfoconcordia* genes in single *O. algarvensis* individuals.** A reanalysis of published metaproteomes showing the top 100 abundant proteins across 10 host individuals (**Table S10**)
